# Lessons learned when looking for non-neutral ecological processes in the built environment: the bacterial and fungal microbiota of shower tiles

**DOI:** 10.1101/413773

**Authors:** Rachel I. Adams, Despoina L. Lymperopoulou

**Author notes:** Corresponding author: Rachel Adams.

## Abstract

With periodic pulses of water, bathroom showers represent a habitat in the built environment with a high potential for microbial growth. We set out to apply a neutral model of microbial community assembly and to identify deviations from the model that would indicate non-neutral dynamics, such as selective pressures for individual taxa, in this particular indoor habitat. Following a cleaning event, the bacterial and fungal microbiota of the shower stalls in two residences in the San Francisco Bay Area were observed over a four-week period. We observed strong differences in composition between houses, preventing us from combining samples and thus limiting our statistical power. We also identified different aspects of the sampling scheme that could be improved, including increasing the sampling area (to ensure sufficient biomass) and increasing the number of replicates within an individual shower. The data from this pilot study indicate that immigrants to the built environment arising from human shedding dominate the shower ecosystem and that growth conditions are relatively unfavorable despite the water availability. We offer suggestions on how to improve the studying and sampling of microbes in indoor environments.

## Introduction

In the built environment of human residences, water is generally a limiting resource. Built to remain dry, most areas of homes are expected to be ecological sinks (as defined by Pulliam 1988), where there is a greater influence of dispersal into the system than present from endogenous growth. An exception to this source-sink framework in homes may be those areas in the bathrooms and kitchens, which receive intentional and frequent water use. These areas where nutrients and periodic pulses of water are available may be ecologically productive, such that microbial communities can become established and interact with each other and their environment; in other words, these wet areas may be true biological ecosystems.

We hypothesized that bathroom showers would be a microbial ecosystem, and so we set out to explore change in the bacterial and fungal communities over time in the built environment. We were inspired by recent work exploring neutral and non-neutral processes that drive microbial assembly in the guts of zebrafish (Burns et al. 2016). As a parallel to zebrafish age, or “days post fertilization”, we considered time since cleaning, which at its best is a process that severely disturbs the established community, eliminating a large portion of existing biomass and creating open niches.

Previous work has shown that both dispersal and selective pressures determine the bacterial composition of bathrooms. Flores et al. (2011) showed that surfaces generally clustered into three types based on their dominant bacterial source populations: surfaces routinely touched with hands, the restroom floor, and toilet surfaces. In a high-occupancy university bathroom setting, the bacterial composition shifted throughout an 8-hour day to one dominated by skin-associated taxa and after this short period, surfaces were compositionally similar to those surfaces that were left for one month prior to sampling (Gibbons et al. 2015). In contrast to surfaces that are exposed to air and human occupants, structures that are part of the plumbing system, such as showerheads and hospital therapy pools, are dominated by a common group of taxa that includes *Mycobacterium, Methylobacterium*, and *Sphingomonas* (Angenent et al. 2005; Feazel et al. 2009; Perkins et al. 2009; Miletto & Lindow 2015). Less work has been done examining fungi on bathroom surfaces, although *Exophiala, Fusarium*, and *Rhodotorula* are common components of drains and dishwashers (Zalar et al. 2011; Adams et al. 2013; ZupanČiČ et al. 2016).

The Sloan neutral model in microbial community predicts the relationship between the occurrence frequency of a taxon in individual local communities and its abundance in the overall metacommunity (Sloan et al. 2006). Using the non-linear least squares as a model of neutral dynamics, one aspect that can be teased out from this application is what taxa deviate from this null model. Those taxa that are more prevalent (have a higher occurrence frequency) than expected based on the model are either being selected for or are frequent immigrants into the system. Those taxa that are less prevalent than predicted based on their overall abundance are either selected against in the system or are dispersal limited.

In the shower tiles of residential bathrooms, the source communities are airborne, waterborne, and human-associated. Once present, the habitat of bathroom tiles has strong selective pressures resulting from relatively unfavorable growth conditions, including desiccation periods and nutrients that are both limited and specific. Nevertheless, even the periodic application of water could create favorable conditions, and we hypothesized that we would be able to, by applying ecological theory, identify those taxa that did not follow neutral processes marked by stochastic loss and replacement of individuals. Ultimately, we discovered that the sampling scheme would have to be highly tailored to this specific environment and proceed for a lengthy duration between cleanings that many would find undesirable for home environments.

## Materials and Methods

### Sample collection

Shower stalls in two houses in the San Francisco Bay Area were sampled for four consecutive weeks in February and March, 2018. Two areas of the shower were targeted, one at the back of the shower opposite the showerhead, and another on the side wall of the shower (Figure 1). Each area was approximately 52 cm wide and 26 cm high and partitioned into four equal size 13 cm x 26 cm subsections. At the start of the sampling period, surfaces were cleaned according to the practices of the house. In House 1, the surface was sprayed with a “disinfecting bathroom bleach-free cleaner”, wiped with a sponge, rinsed with water, dried out with a washcloth, and finally cleaned with an ethanol wipe. In House 2, the surface was sprayed with a “naturally derived tub and tile cleaner” and washed with a washcloth for approximately 30 seconds.

Approximately one week after cleaning, the first partition was sampled by rubbing a water-moistened Floq swab back-and-forth over the first subsection area for approximately one minute. The swab tip was broken off into an Eppendorf tube and immediately frozen.

Approximately two weeks after cleaning, one swab was used to sample the subsection that had been sampled the previous week and a second swab was used to sample the next subsection that had been left undisturbed since cleaning. This continued for four weeks and resulted in 10 swabs for a given area of the shower stall, either back or side wall. Sampling was approved by the University of California Committee for the Protection of Human Subjects under protocol 2015-02-7135.

**Figure 1.**
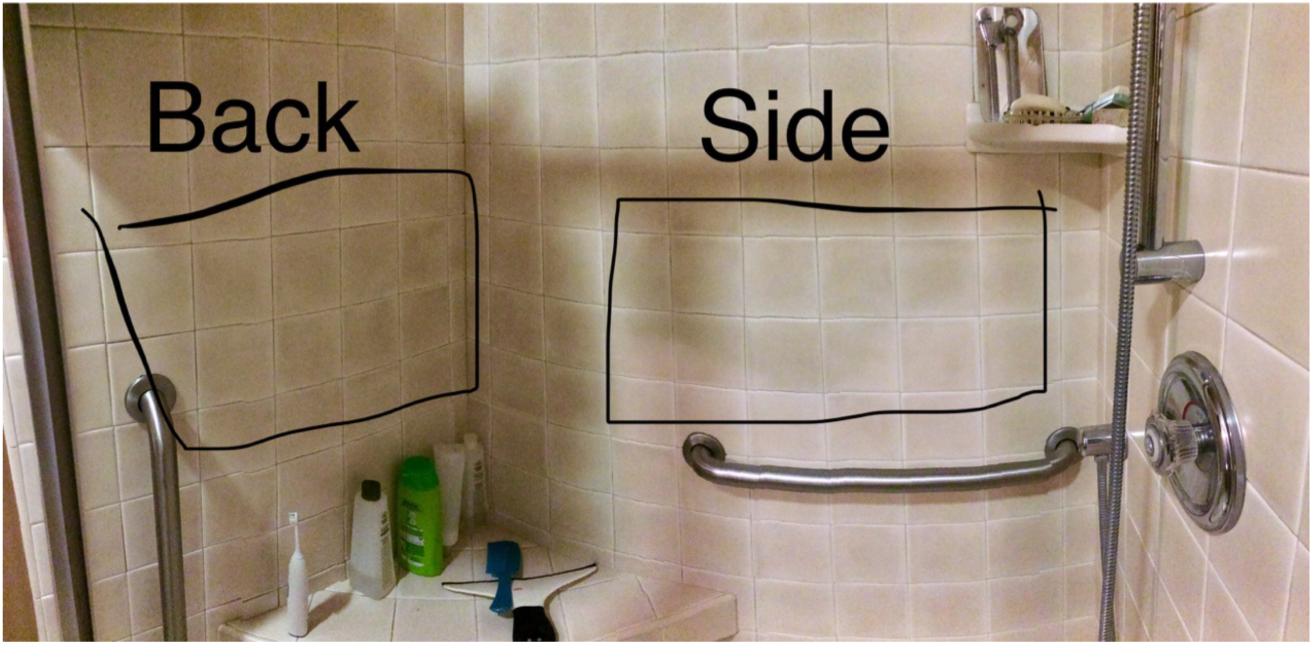
Image of one of the shower stalls studied, highlighting that two different areas of the stall were targeted.

### Sequence generation and analysis

Genomic material in swabs were extracted using the Qiagen PowerSoil Kit and followed the manufacturer protocol. Libraries were prepared at the QB3 Vincent J. Coates Genomics Sequencing Laboratory at the University of California, Berkeley, using a two-step PCR process (Detailed methods are included in Supplementary File 1). For bacteria, the 16S v4 hypervariable gene region was targeted using the 515f forward primer and 806r reverse primer in the first PCR. For fungi, the first PCR targeted the ITS1 region of the 18S gene using the ITS1f forward primer and ITS2 reverse primer. The second PCR reaction adds Illumina adapters and barcodes to either side of the PCR products. Both positive and negative samples were included. For bacteria, we purchased a mock microbial community standard (Zymo Research). For fungi, non-biological synthetic mocks were cloned and combined into a community (Palmer et al. 2018).

Extraction and PCR negatives were also processed along with our biological samples. All the samples were pooled and run on a single 300PE v3 MiSeq lane. Sequences have been deposited in NCBI’s Sequence Read Archive (SRA) with accession SRP148593.

Bacteria sequenced were processed using the DADA2 pipeline (Callahan et al. 2016) implemented in R (R Development Core Team 2014). Detailed commands are included in the Supplementary file and briefly discussed here. Forward and reverse reads were filtered and trimmed, and the error rates learned. Following dereplication, amplicon sequence variants (ASVs) were inferred on the pooled data. Forward and reverse reads were then paired, and a table was constructed from the merged reads. Chimeras were removed, and taxonomy assigned against the SILVA database (Quast et al. 2013).

Fungal sequences were processed using AMPtk v1.0.2 (Palmer et al. 2018). Reads were preprocessed, specifying the ITS1-F and ITS2 primers were not requiring. This program relies on internal scripts as well as usearch (Edgar 2010) and vsearch (Rognes et al. 2016) to merge paired reads, remove PhiX, quality filter, trim any remaining primer sequence, cluster sequences, and remove chimeras. Taxonomy was assigned using a combination of sintax (Edgar 2016) and utax against the UNITE database (28.06.2017) (Koljalg et al. 2005), with the sequences of the synthetic mock added. Detailed commands are included in the Supplementary file.

Community tables were analyzed using the Phyloseq package (McMurdie & Holmes 2013) in R (R Development Core Team 2014). Contaminant sequences were identified using the decontam package (Davis et al. 2017) at the 50% prevalence threshold for bacteria and 30% threshold for fungi. The program identified 112 bacterial contaminants and 21 fungal contaminants, and these were removed from their respective community tables. Taxa that were classified as Eukaryota (n = 83) or left unclassified (Phylum=NA, n=34) were removed from the bacteria community table. Assessment of the performance of the positive mocks were evaluated following these filtering steps. For the bacteria mock, the eight bacteria taxa were recovered at high sequence read count (>10,000 sequences), with an additional 52 taxa represented at lower sequence count (< 2,600). Most of these were not present in study samples, and, based on their shared taxonomic identity with those in the mock, likely resulted from imprecise grouping of mock sequences. With the fungal mock, the eight synthetic mock sequences were recovered at high sequence read count (> 25,000 sequences), with an additional seven taxa represented at low sequences abundance (< 11 reads).

## Results and Discussion

We frame our results in the form of main findings and lessons learned in order to inform future efforts when looking for non-neutral ecological processes in built environments.

### 1. Houses are not zebrafish

We approached this work viewing houses as technical replicates, as was done in previous work where separate buildings are sampled in order to identify patterns common across the different settings (e.g. Flores et al. 2011; Dunn et al. 2013; Gibbons et al. 2015). We observed strong differences in the microbial communities across these two houses, particularly for fungi (Figure 2). House 2 was dominated by two fungi, *Didymella glomerata* and *Filobasidium magnum*, and this stayed constant throughout the sampling period. In contrast, the fungal community in House 1 was much more diverse (Shannon diversity, mean House 1 = 2.5; mean House 2 = 0.64; t-test p<0.001). Using the Bray distance index, house alone explained 44% of the variance between fungal taxa (adonis p<0.001). Bacteria showed a similar pattern of lower diversity in House 2, but the difference was less extreme (Shannon diversity, mean House 1 = 4.3; mean House 2 = 3.7; t-test p<0.001). Again, using the Bray distance index, house explained 30% of the variance between bacterial taxa (adonis p<0.001). The strength of the house effects are consistent with another study looking at the bacteria on kitchen sinks (Moen et al. 2016).

The strong microbial communities between houses required that investigations in neutral and non-neutral processes occured within an individual house. That is, unlike zebrafish, the samples from different houses could not be combined into a common dataset. This strong house-specific effect is not altogether surprising, given the many ecological factors that could bring about this pattern, notably founder effects and different source populations arising from geographic and plumbing system variation. Nevertheless, efforts to characterize the microbiota in buildings typically draw on many different houses. For example, the 1000 Homes Project was a community-based sampling effort across the entire United States; it showed that the bacteria in house dust were largely determined by occupants and their activities, while fungi were largely determined by outdoor environmental characteristics (Barberán et al. 2015a; Barberán et al. 2015b; Grantham et al. 2015). On the other hand, a study looking at the interactions between human occupants and dry surfaces in indoor environments (such as doorknobs, light switches, and floors) focused on repeated sampling of six homes (Lax et al. 2014). A larger version of this pilot study would benefit from this second kind of sampling scheme, in that it balances representation across homes with sufficient sampling depth to investigation temporal dynamics.

**Figure 2.**
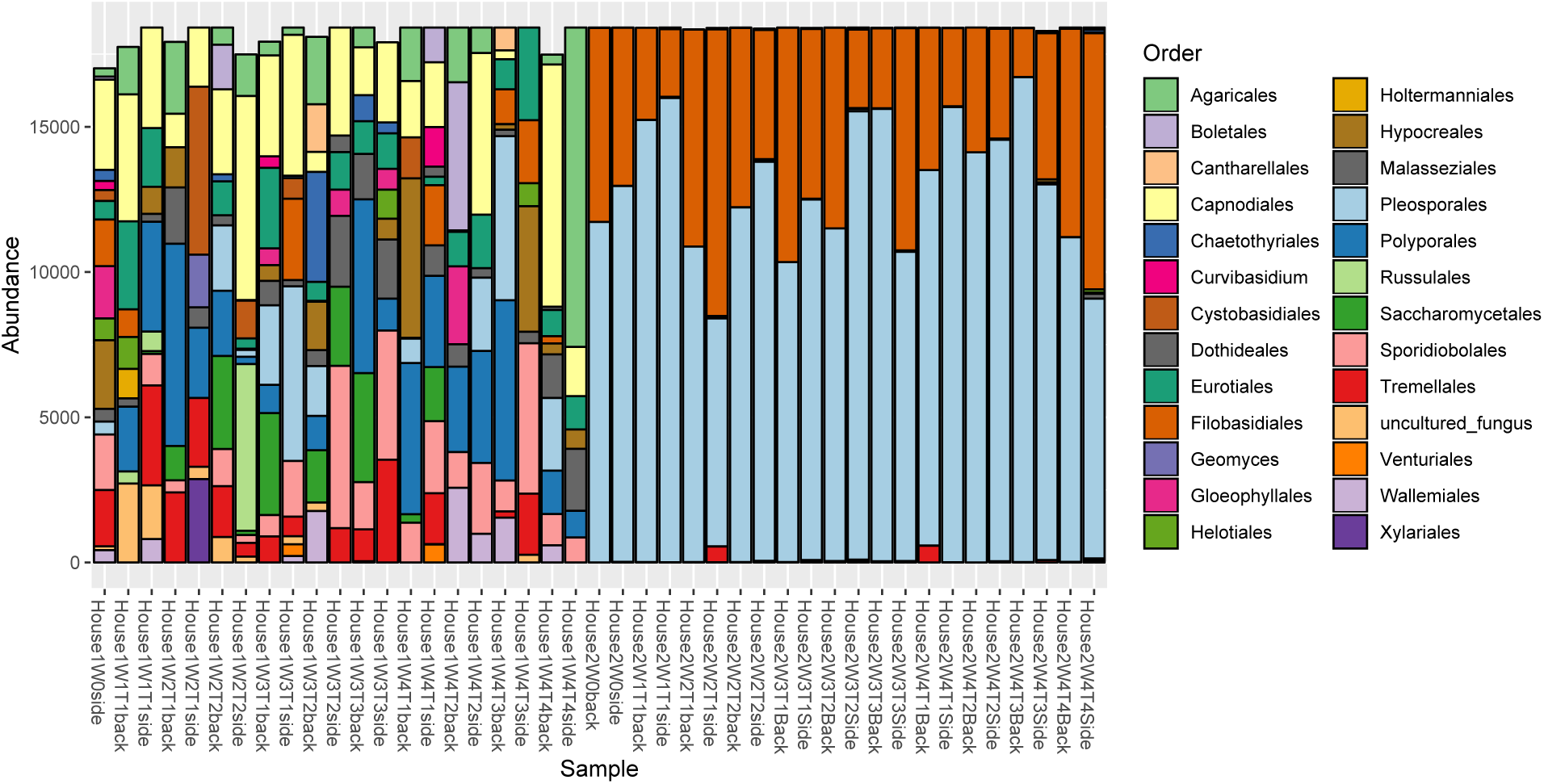
Overview of fungal orders between the two houses. Samples are named according to house (House 1 or House 2), the week it was sampled (W0-W4), the tile that was sampled (T1-T4), and the location in the shower (back or side).

### 2. Identifying representative sampling

Spatial variation within a specific surface type in the home environment has not been systematically studied. Spatial structure could arise if the orientation of the stall and occupant create differential source strengths of the water from the premise plumbing system and what splashes off the occupants. In a previous study, we found that variation in community composition across two different areas of a kitchen sink and a shower stall was less than the variation observed across homes (Adams et al. 2017). In agreement with our previous observation, we found that, within a house, there was little or no impact of location (back or side) within the shower on fungal composition (adonis, House 1: R^2^=0.07, p=0.01; House 2: R^2^=0.003, p=0.87), but a stronger effect of location for the bacterial composition (adonis, House 1: R^2^=0.13, p=0.01; House 2: R^2^=0.21, p=0.001). Given these results, the value of replicates exceeds the potential information gleaned from spatial structure. Thus, many different areas of the shower stall should be sampled at a given time point, and these can serve as time point replicates in the data analysis stage.

Swabbing surfaces should represent a disturbance to the microbiota assembled there, so we sampled a new tile each week as well as the tiles sampled previously (for example, in week 3, we sampled tile 3, as well as took repeat swabs of tiles 1 and 2 that had been swabbed in previous weeks). In the analysis, these samples of re-swabbed areas provided little, if any, information on the dynamics of the system. The dominant taxa remained consistent across the weeks, regardless of how many times that area had been swabbed, and it was low abundant taxa that differed across weeks. Swabbing was just one of many factors that could have caused the fluctuation of low abundant taxa. Other factors that could affect low abundant taxa include changes in the waterborne taxa introduced into the shower and stochastic splashing off of the shower occupants. Our study was not able to tease out the specific effect of swabbing or these other factors. Thus, we conclude that resampling previously swabbed areas can be skipped in future sampling efforts.

Sufficient biomass collection in indoor environments is a long-standing problem. While some studies report community profiles from what seems like small biomass, such as from keyboards (Fierer et al. 2010) and light switches (Lax et al. 2014), we found that the yield from the area we sampled was approaching the detection limits of our approach. The reads counts of some of the samples approximated those of some of the negative controls (Figure 3). Recent work shows that including more precise positive controls can help with interpretation of these kinds of low biomass samples (Minich et al. 2018).

**Figure 3.**
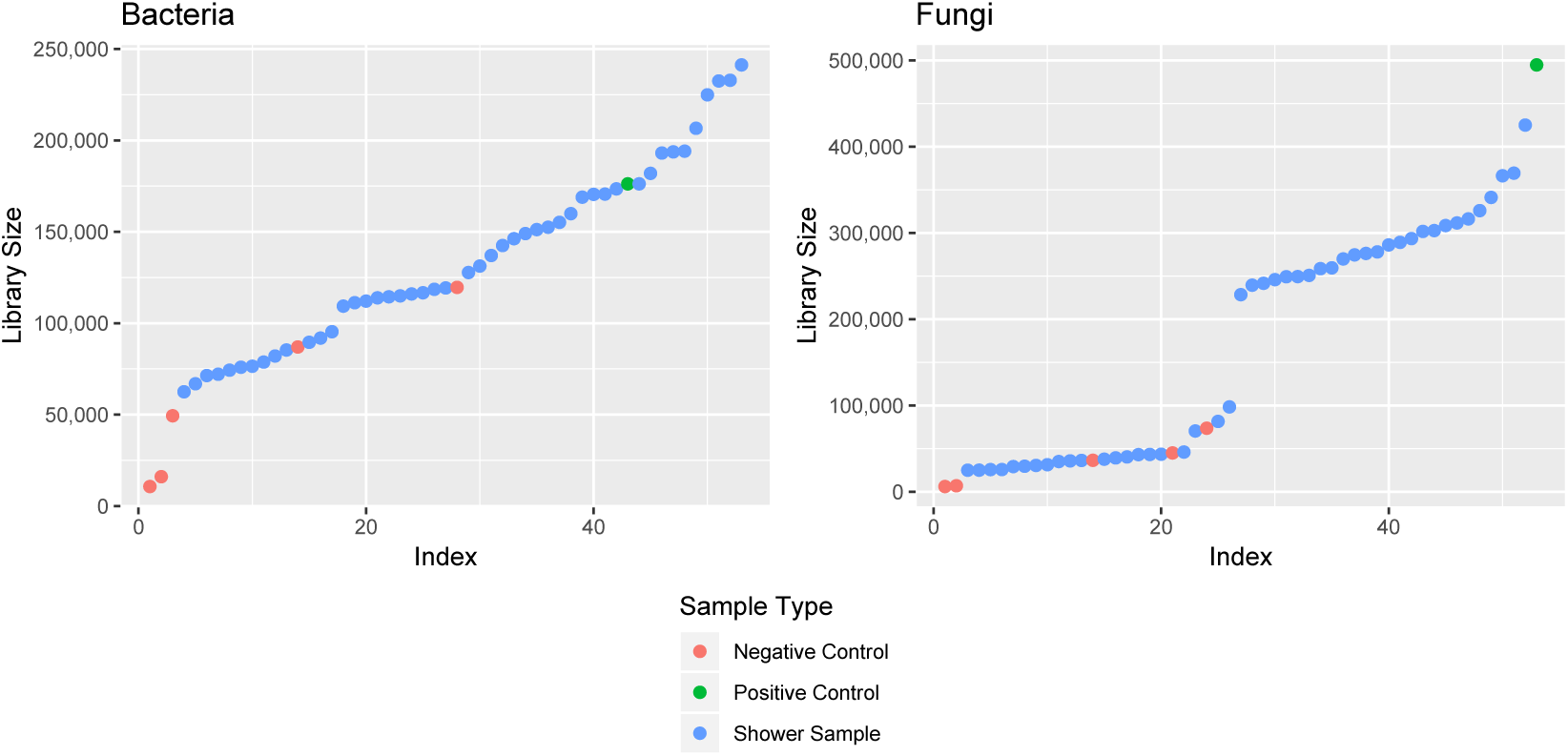
Sequence counts of bacterial and fungal samples following removal of taxa using the “decontam” package showing that biomass for many shower samples approximate those of some of the negative controls.

## 3. Results of Sloan model and study duration

Due to our limited sample size, results from applying a neutral model of community dynamics should be taken with caution. While we had eight samples for the week 4 time point, Burns et al. used approximately 20 fish for each time point. Many bacteria were less prevalent than predicted based on their abundance (bottom right areas in Figure 4); these are either selected against, or are dispersal limited system. In both houses, partitions below the neutral prediction were most strongly distinguished by the presence of Proteobacteria and Firmicutes, in particular the family Sphingomonadaceae and order Clostridiales. There were many more bacteria that were more prevalent than predicted by the model (top left in Figure 4); these are either selected for or are frequent immigrants into the system. Partitions above the neutral prediction were distinguished by Actinobacteria and to a lesser extent Proteobacteria, and were particularly represented by Staphylococcaceae, Streptococcaceae, and Corynebacteriacea (taxa are commonly associated with human skin (Byrd et al. 2018)), and by Mycobacteriaceae and Methylobacteriaceae (taxa associated with premise plumbing (Falkinham et al. 2015)).

Similarly, skin-associated taxa were the most common type of bacteria to increase in abundance over the sampling period (Figure 5).

**Figure 4.**
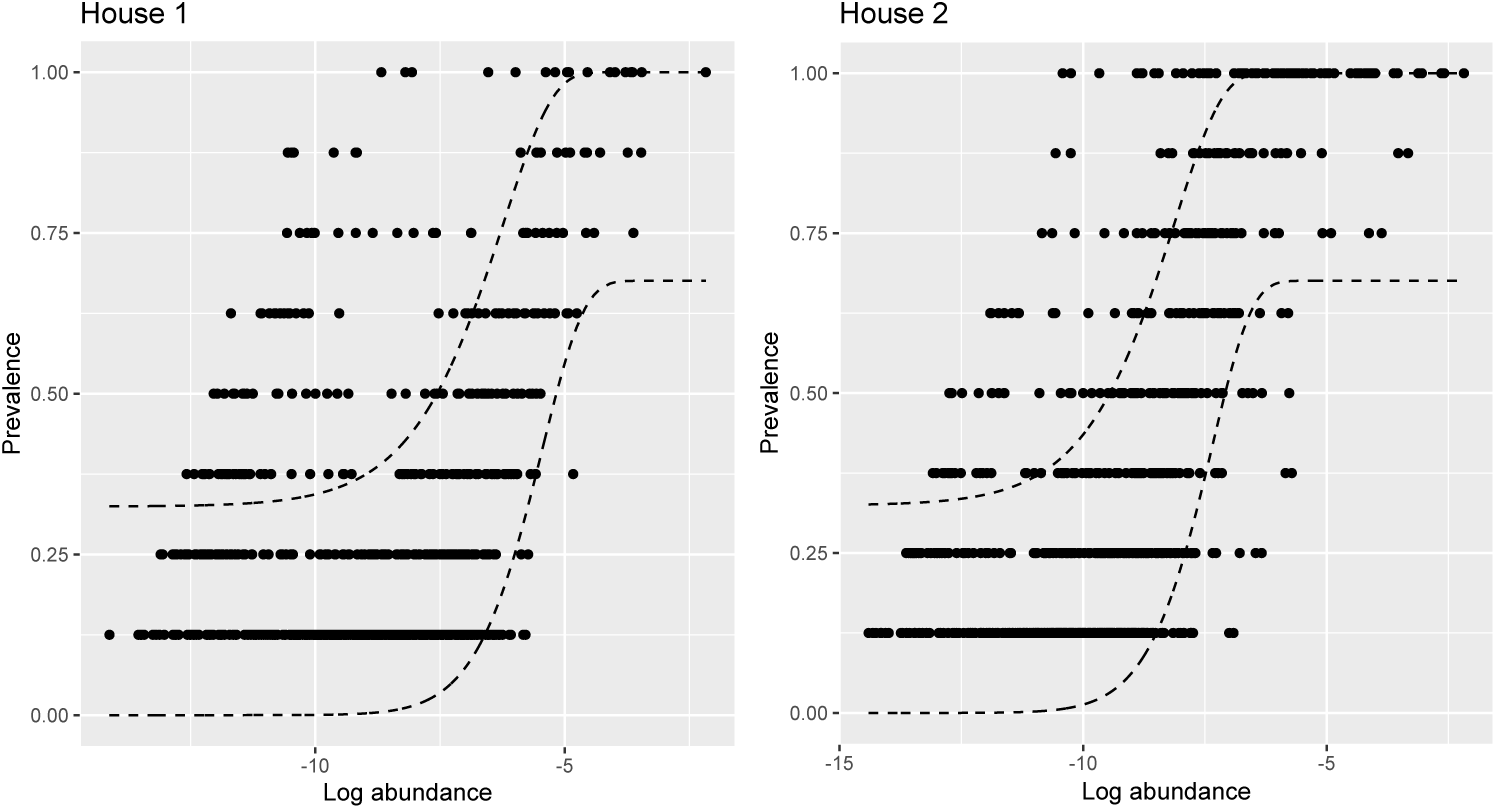
Individual ASVs of bacteria in the 4^th^ week of sampling and their fit with the Sloan neutral model.

**Figure 5.**
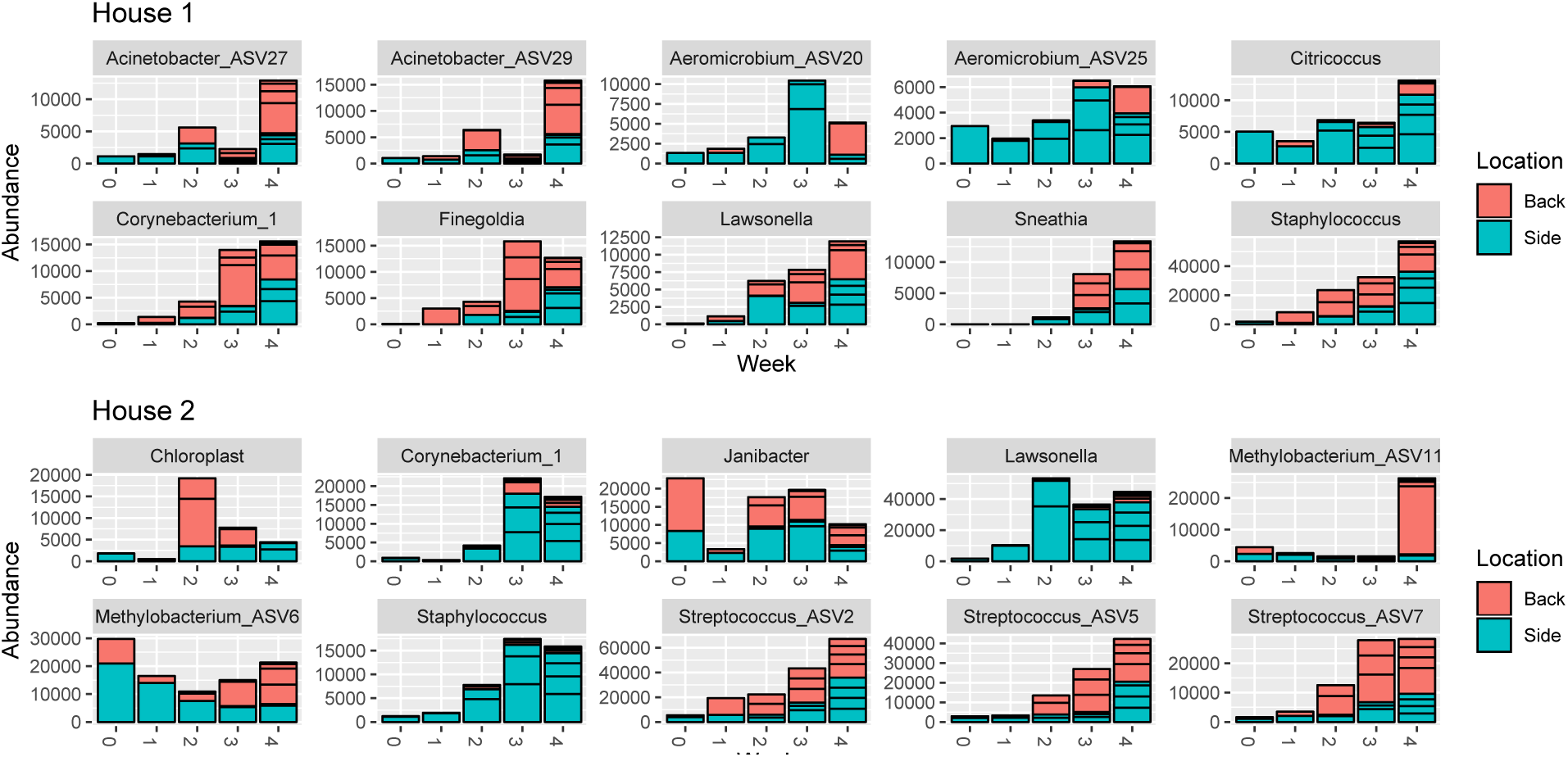
The change in abundance of the top most abundant bacteria ASVs across the sampling period.

There were many fewer fungal taxa flagged as deviating from the null model in household showers (Figure 6). Taxa lower than predicted (i.e. selected against or dispersal limited) were frequently taxa associated with outdoor environments, including plant pathogens (*Gibberella baccata* (Afanide et al. 1976) in House 2) and soil fungi (*Fusarium oxysporum* (Michielse & Rep 2009) and *Papiliotrema terrestris* (Liu et al. 2015) in House 1). Fungal taxa that were more prevalent that predicted by the model, based on their abundance, were overwhelmingly members of the Malasseziales order, indicating that this skin-associated fungus (Findley et al. 2013) is a frequent immigrant into this system. Accordingly, the abundance of Malassezia does increase over time (Figure 7). Other fungi, such as *Filobasidium, Rhodotorula, Cladosporium*, and *Papiliotrema*, also show increasing abundance over time, highlight which fungal taxa are likely to become increasingly important components of the system should the environment be left undisturbed by cleaning.

**Figure 6.**
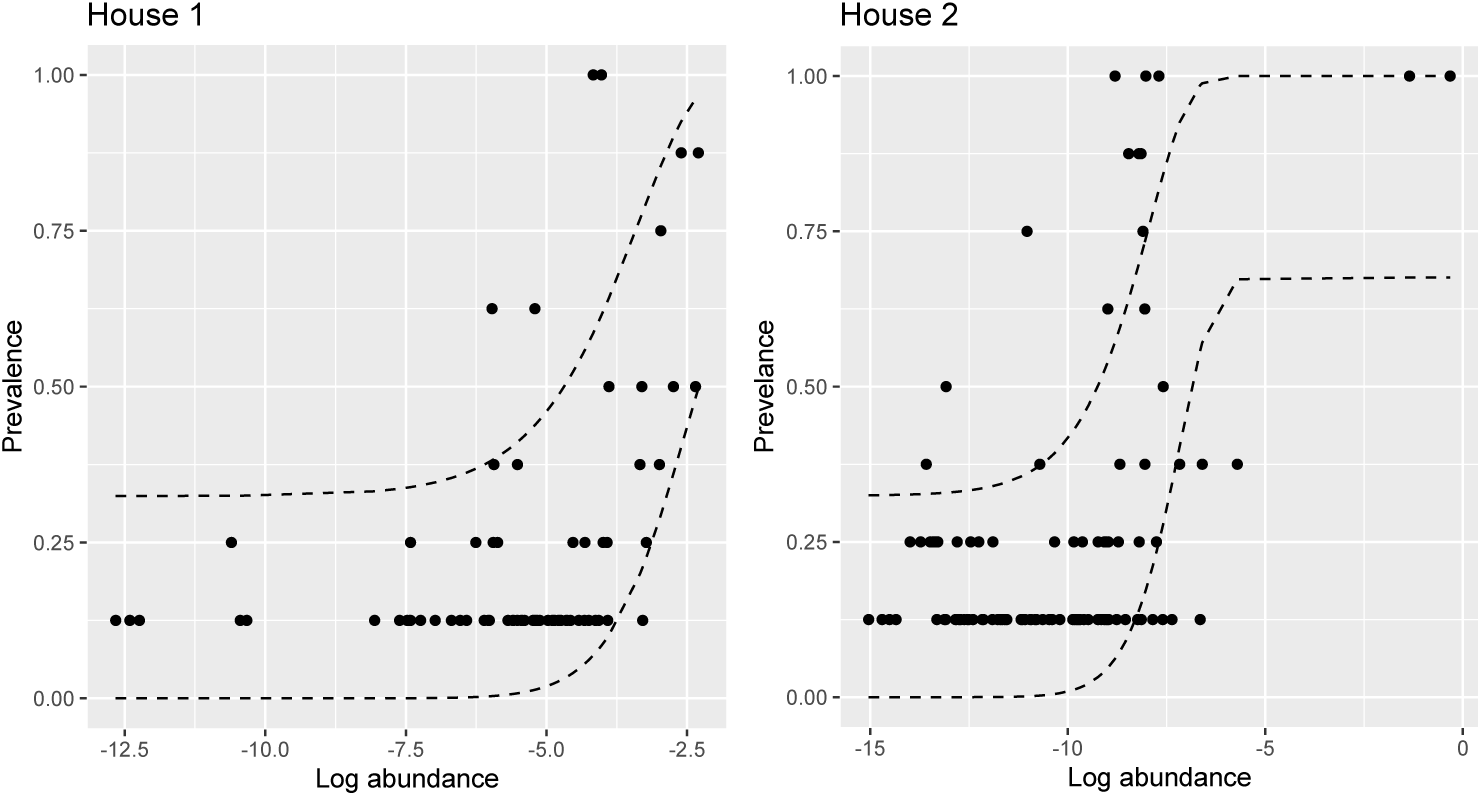
Individual OTUs of fungi in the 4^th^ week of sampling and their fit with the Sloan neutral model.

**Figure 7.**
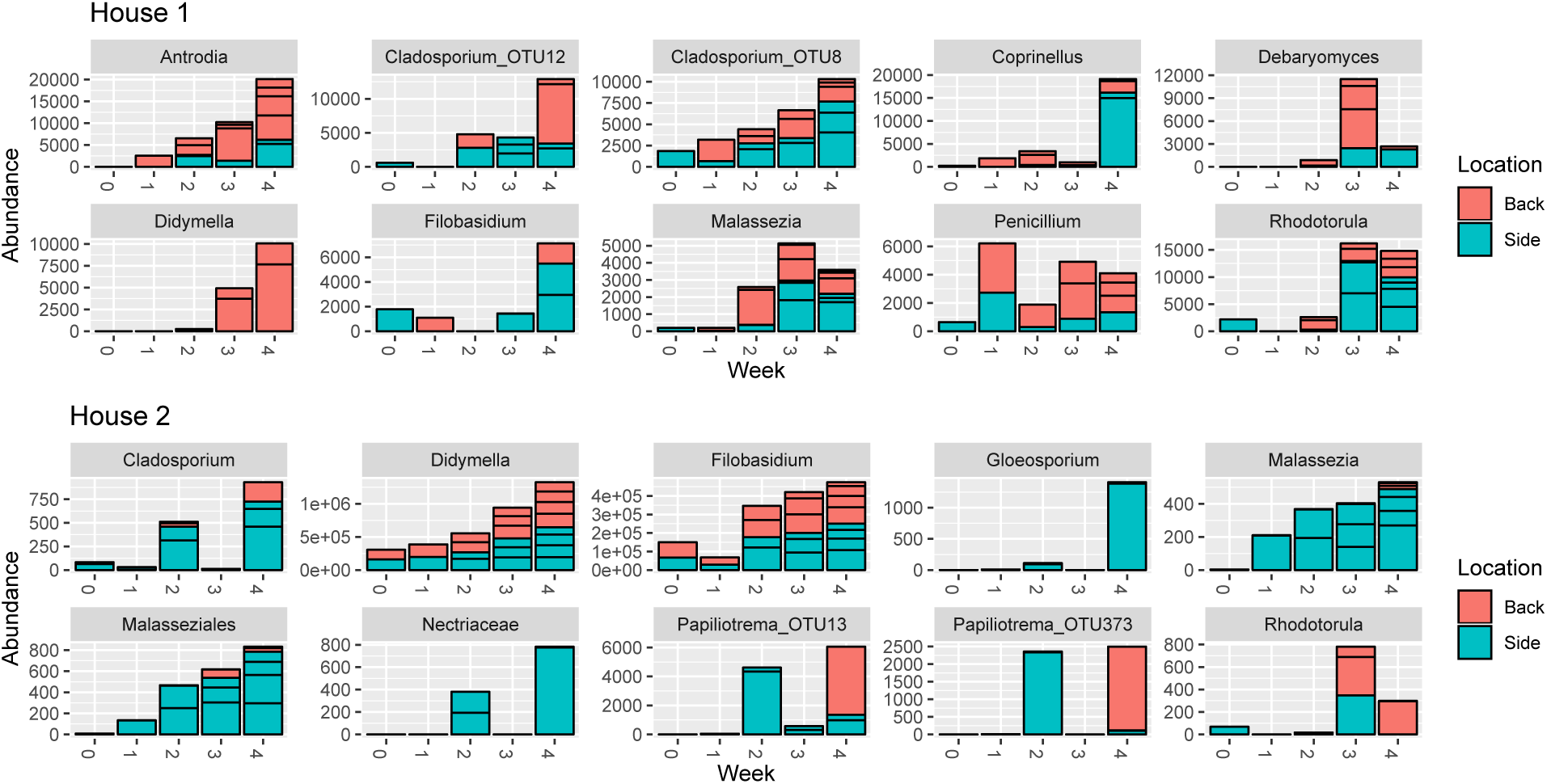
The change in abundance of the top most abundant fungal OTUs across the sampling period.

## Conclusions

Given all these considerations, we would make the following recommendations as a sampling scheme for longitudinal sampling of shower stalls.

- Inclusion of more than two houses is necessary to ensure a representative sample of houses.
- At least 15 areas of the shower stall should be sampled at a given time point. These samples will represent different samples in the larger metacommunity of that time point.
- The sampling area within the shower should be large vertical areas, extending as large as possible to balance sufficient surface area to swab with enough area in the shower to allow for undisturbed areas to swab in subsequent weeks. We anticipate that an area of 10cm x 100cm would yield sufficient biomass and allow room for repeated sampling over weeks. In subsequent sampling periods, the sampling area could shift horizontally.
- Sampling should take place every two weeks and continue for at least ten weeks post cleaning.

Despite the technological hurdles of implementation, these preliminary data indicate that, even in areas of the household environment that are relatively favorable for microbial growth compared to other (i.e. drier) places, immigrants arriving from human shedding dominate the microbiota and that growth conditions are relatively still unfavorable for the time duration we examined. It is unclear whether these taxa take up residence on bathroom showers or simply persist on skin fragments and accumulate over time, and other tools, such as using isotopically-labelled compounds (e.g. Berry et al. 2015), may be a more powerful approach to identifying true ecological residents. These data also show that after a 4-week period, evidence for deterministic processes is limited, indicating that periodic cleaning of at least this frequency is sufficient to keep microbial communities associated with bathroom tiles in a non-steady state of limited productivity.

## Acknowledgements

We thank Annie Lai for help with sample processing, Dylan Smith for library preparation, and Jon Palmer for assistance with running AMPtk.

